# A physiological microfluidic blood-brain-barrier model for in vitro study of nanoparticle trafficking and accumulation

**DOI:** 10.1101/2025.08.28.672885

**Authors:** Bryan B. Nguyen, Neona M. Lowe, Sophia Kellogg, Kuan-Wei Huang, Hannah O’Toole, Elizabeth J. Hale, Venktesh S. Shirure, Bhupinder S. Shergill, Steven C. George, Randy P. Carney

## Abstract

Although the blood-brain barrier (BBB) restricts passage of most molecules, various naturally occurring and synthetic nanoparticles are nonetheless found within the brain parenchyma. To study the mechanisms underlying this phenomenon, we developed a microfluidic BBB model (mBBB) using human cerebral microvascular endothelial cells (HCMECs) in direct contact with primary human astrocytes and pericytes within a physiologically relevant extracellular matrix. The horizontal architecture enables high-resolution imaging across the full barrier interface and allows direct assessment of nanoparticle transport and accumulation. This *in vitro* platform recapitulates key features of the BBB, including selective permeability, junctional protein expression, and receptor-mediated uptake pathways. Using this system, the trafficking and accumulation of structurally distinct nanoparticles, including liposomes, nanoplastics, and extracellular vesicles (EVs), were compared. Among these, heterologous EVs exhibit the highest transport efficiency. Analysis of nanoparticle properties suggest that ligand presentation and membrane composition, rather than size or stiffness, primarily govern BBB penetration. The mBBB platform provides a high-throughput, imaging-based framework to systematically interrogate nanoparticle trafficking across the BBB and offers a translational tool for both drug delivery and neurotoxicity screening.

## 1. Introduction

The blood-brain barrier (BBB) is a highly regulated structure comprised of endothelial cells, pericytes, and astrocytes.^1^ Upregulated adherent and tight junction proteins on the endothelial component of the BBB, which are further reinforced via supporting pericytes and astrocytic feet, work together to severely restrict the transport of biomolecules and particles larger than a few nanometers.^2^ The BBB endothelium also expresses a variety of specialized transporters that regulate the transcellular passage of nutrients, waste, and exogenous materials.^3^

Paradoxically, larger nanoparticles with an average diameter between 10 to 100 times the paracellular width (≲5 nm),^4^ can still be found in the brain parenchyma behind the intact BBB.^2,5^ This observation has been highlighted for both its potential for therapeutic development as well as its growing environmental health concern.^6,7^ Over the past few decades, significant research efforts have been focused on understanding and exploiting naturally occurring and synthetic nanoparticles for BBB penetration.

Extracellular vesicles (EVs), heterogenous lipid-bilayer encapsulated nanoparticles produced by all cells, play a critical role in cell-to-cell communication. EVs derived from cells outside the brain parenchyma have been found in the brain, contributing in part to dysfunction and pre-metastatic niche formation in cancer.^8^ Similarly, environmentally generated nanoplastics, small particles resulting from the degradation of plastic products or pollution, can accumulate in brain tissue and are linked to various neurological diseases.^6,9^ Synthetic nanoparticles have also been explored due to the inherent benefit of their scalability and modularity, and have been designed to take advantage of certain characteristics of natural nanoparticles to facilitate transport across the BBB.^10^ Characteristics such as size, charge, stiffness, composition, and ligand presentation have all been tuned to mediate BBB penetration.^7,11,12^

While increasing numbers of brain-penetrant NP formulations have emerged, a commensurate increase in FDA-approved brain-penetrant therapeutics has not.^13^ This gap is exacerbated by the lack of existing *in vitro* models that can replicate key compositional and spatial characteristics of the BBB, and thus are incapable of providing mechanistic insight into how nanoparticles traffic and accumulate.

Such mechanistic insight is essential to advance therapeutic innovation and public health protection. For nanotherapeutics, uncovering the features that enable efficient, safe, and reproducible BBB penetration will guide the rational design or selection of nanoparticles (e.g., EV subpopulations) as therapeutic delivery vehicles to treat neurological diseases. On the safety front, given increasing evidence linking certain nanoparticles to neurodegenerative disorders and cancer, understanding entry and accumulation mechanisms is crucial to identify risk factors and design mitigation strategies.

Finally, such insights better enable effective predictive modeling (e.g., for FDA guidance) to streamline screening and regulatory approval for therapeutic candidates.^14–19^

Currently, no complex multicellular model has been used to systematically compare the different types of NPs and how differences in structural and biochemical features may dictate both access across and penetration beyond the BBB into brain parenchyma. With this goal in mind, we developed a novel *in vitro* microfluidic blood-brain barrier (mBBB) platform that recapitulates the human BBB’s three-dimensional multicellular architecture.

Our mBBB model features a spatially defined co-culture of endothelial cells, pericytes, and astrocytes embedded in a tunable extracellular matrix (ECM). The cells and matrix are arranged in a horizontal, parallel-channel microfluidic design that enables direct visualization of nanoparticle penetration and cell-specific interactions across the barrier. Unlike traditional Transwells^TM^ or spheroids that only exhibit a few of the following features, our mBBB supports dynamic flow across the endothelium, interstitial flow through the endothelium and matrix, facilitates high-resolution long-term imaging, allows for recovery of nanoparticles that cross the BBB, and can be perturbed by chemical treatments to investigate various mechanisms of entry. Critically, the model is constructed using standard soft lithography, and is seeded using routine cell culture techniques, thus encouraging reproducibility and scalability. Its modularity permits future integration of additional cell types or disease-relevant features, offering a flexible platform for both mechanistic studies and translational screening.

Here we apply this mBBB model to interrogate how structurally and biochemically distinct nanoparticles, including synthetic liposomes, biologically derived EVs, and environmentally relevant nanoplastics, interact with and traverse the BBB. The model’s unique horizontal channel architecture allows direct visualization of nanoparticles across the BBB, enabling measurement of penetration depth, cellular localization, transport efficiency, and extent of modification during traversal. By systematically comparing diverse nanoparticle classes, the mBBB can uncover whether trafficking is governed by shared or distinct mechanistic pathways, driven by combinations of particle size, structure, and biochemical composition. These studies address a critical gap: we currently lack a reliable *in vitro* framework linking nanoparticle properties to specific entry routes. Our findings lay the foundation for a new conceptual model of BBB trafficking mechanisms and pose a central question: how do different nanoparticles gain access to the brain, and what mechanistic rules underlie this selectivity?

## 2. Results

### 2.1 Establishment and validation of a microfluidic model of the BBB to study nanoparticle transport

The mBBB device is comprised of five parallel channels separated by microposts, enabling spatial segregation of endothelial cells, astrocytes, and pericytes, while permitting crosstalk at defined interfaces (**Fig. 1A**). The dimensions and hydrostatic pressure lines mimic *in vivo* flow and shear in the brain through microvessels to facilitate BBB development (Fig. 1B).^20^ The central “vascular” channel is seeded with human cerebral microvascular endothelial cells (HCMEC/D3), chosen due to its well-documented low permeability and physiologic relevance,^21,22^ while adjacent compartments support primary human astrocytes and pericytes embedded in a fibrin ECM. The horizontal configuration allows direct imaging of nanoparticle distribution across the BBB layers under physiologically relevant conditions, including intermittent flow and shear, which are known to improve endothelial phenotype and facilitate supporting cell interactions.^20,23–27^

**Figure 1.**
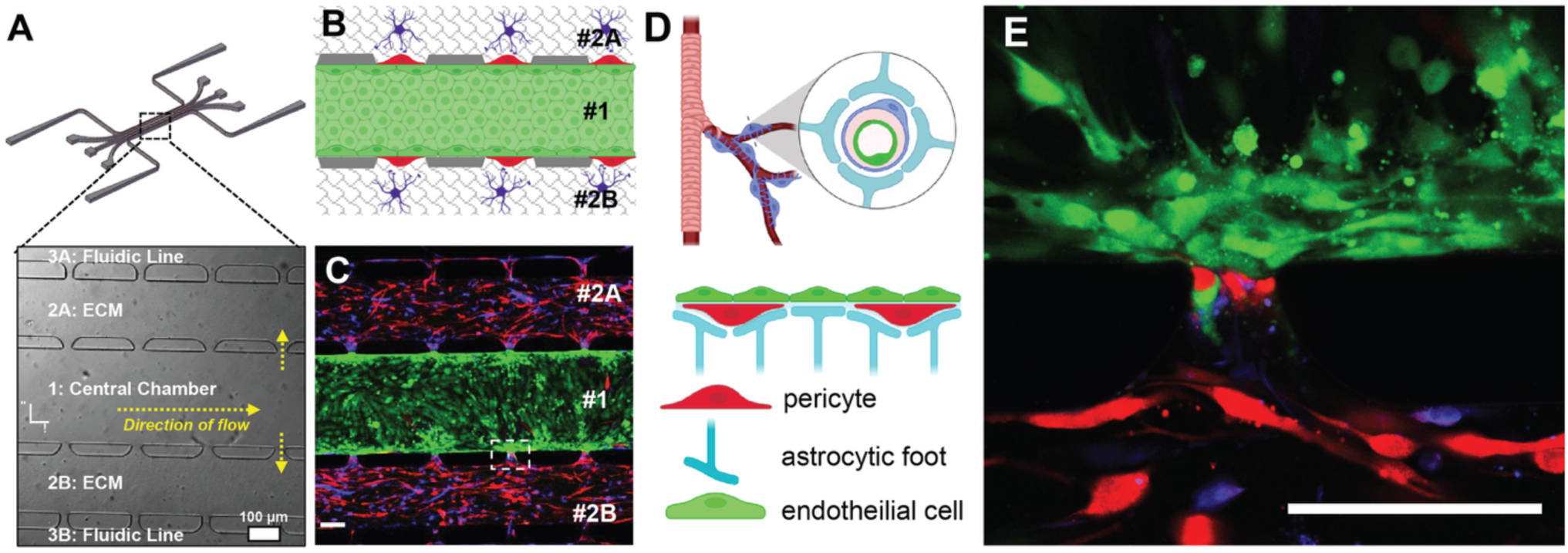
Development and characterization of a physiologically structured microfluidic BBB model (mBBB). (**A**) Overview of the microfluidic device used to establish the mBBB. Bottom: Brightfield image showing five parallel channels that communicate through a series of pores ∼30 μm in diameter. The central channel (#1) is seeded with endothelial cells (ECs), while adjacent ECM channels (#2A, #2B) contain astrocytes and pericytes embedded in fibrin matrix. Yellow arrows indicate flow direction and crosstalk interfaces. **(B)** Schematic cross-section of the mBBB, illustrating tri-cellular architecture with ECs (green) forming a continuous monolayer in the central channel, flanked by astrocytes (blue) and pericytes (red) in the ECM channels. **(C)** Confocal image of the fully formed mBBB, showing distinct layers of endothelial cells (green), pericytes (red), and astrocytes (blue) aligned across the communicating pores. Magnification: 10x. **(D)** Top: Anatomical context of the in vivo neurovascular unit, highlighting pericyte and astrocytic interactions with the vascular wall. Bottom: Simplified schematic of tri-cellular interaction in the mBBB, showing pericyte-endothelial and astrocyte-endothelial interfaces. Image generated in BioRender. **(E)** High-magnification confocal image of the interface zone (pore) showing direct contact between endothelial cells, pericytes, and astrocytic processes, recapitulating BBB-relevant spatial relationships. Magnification: 60x. For all images, laser: 488 nm, 561 nm, 640 nm; gain: 1.0; Exposure time: 2 µs/pixel: scale bar: 100 µm.

To verify the structural integrity and cellular architecture of the mBBB, we performed confocal microscopy following fluorescent cytosolic labeling. We observed confluent endothelial layering in the central channel and clear alignment of the three cell types at the micropore interfaces, indicating effective tri-cellular organization (**Fig. 1B-E**). This configuration enables discrete barrier formation at the interface zones where all three cell types interact, mimicking the *in vivo* spatial microenvironment of the BBB.

We next assessed whether the mBBB recapitulates hallmark features of *the in vivo* BBB. Immunofluorescence staining revealed robust expression of tight junction proteins (ZO-1), gap junction components (connexin-43),^28,29^ and multiple key transport and efflux proteins, including transferrin receptor (TfR),^30,31^ LDL receptor (LDLR), scavenger receptor class B type I (SR-BI),^32^ and P-glycoprotein (P-gp)^33^ (**Fig. 2A**). While expression of claudin-5 was relatively weak, consistent with known limitations of the HCMEC/D3 line,^34^ the overall pattern of protein expression supports the phenotype of the BBB.

**Figure 2.**
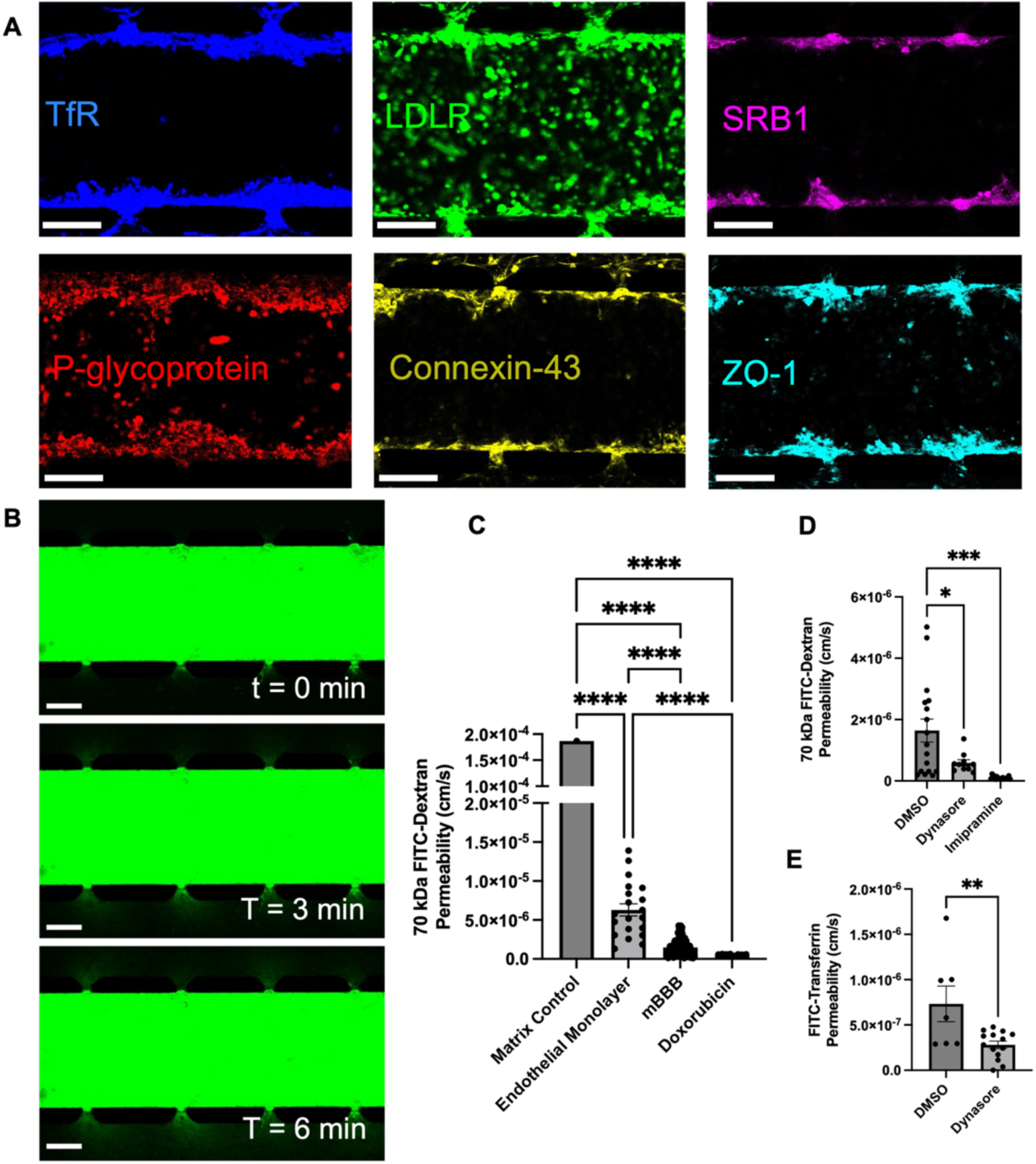
The mBBB recapitulates key functional and molecular characteristics of the in vivo BBB. **(A)** Immunofluorescence staining of the mBBB showing expression of canonical tight junction and transporter proteins, including transferrin receptor (TfR), LDL receptor (LDLR), scavenger receptor class B type I (SR-BI), P-glycoprotein, Connexin-43, and ZO-1. Magnification: 10x. Laser: 640 nm. Gain: 1.0. Exposure time: 2 µs/pixel. Scale bar: 100 µm**. (B)** Time-lapse imaging of 70 kDa FITC-dextran diffusion from the vascular channel into adjacent ECM channels over a 6-minute interval, showing restricted permeability. Magnification: 10x. Laser: 488 nm. Gain: 1.0. Exposure time: 2 µs/pixel. Scale bar: 100 µm**. (C)** Quantified 70 kDa dextran and doxorubicin permeability across control (matrix only n=1), endothelial monolayer (n=20), and mBBB (n=165). Permeability is significantly reduced in the mBBB relative to controls by one-way ANOVA. Doxorubicin permeability (n=8) across the mBBB is lower than for dextran, consistent with active efflux via P-glycoprotein by one-way ANOVA. **(D)** Both pharmacological inhibition of dynamin-mediated endocytosis with Dynasore (n=10) and inhibition of macropinocytosis with Imipramine (n=17) significantly reduce FITC-Dextran permeability when compared to control (n=17), indicating the mBBB supports multiple uptake pathways that can be modulated pharmacologically. **(E)** Dynasore (n=14) significantly decreases FITC-Transferrin permeability when compared to control (n=7), confirming functional receptor-mediated transport by unpaired t-test. (*) indicates p < 0.05, (**) indicates p < 0.01, (***) indicates p < 0.001, (****) indicates p < 0.0001, and (ns) indicates p > 0.05. Error bars indicated standard error in the mean (SEM). Replicate data are biological replicates. Sample size (n) indicates pores used in analyses.

To determine the functional permeability characteristics of the mBBB, we quantified transport of 70 kDa FITC-dextran and doxorubicin under standard culture conditions. Diffusion of FITC-dextran from the central channel to the adjacent ECM-filled compartments was minimal over a 5-minute time course, yielding an average permeability coefficient of 1.47 × 10^-^⁶ cm/s (**Fig. 2B-C**). This is markedly lower than values measured in endothelial monocultures and is consistent with previously reported *in vitro* BBB models.^35,36^ Doxorubicin, a chemotherapeutic known to be effluxed by P-gp,^37,38^ and with a molecular weight (543.5 Da) similar to the dextran, exhibited even lower permeability across the mBBB (5.7 × 10^-^⁷ cm/s) similar to previously reported *in vivo* measurements,^39–41^ further supporting functional transporter activity (**Fig. 2C**).

Finally, we pharmacologically perturbed two canonical endocytic pathways to test whether known mechanisms of BBB transport are present and active. Pre-treatment with Dynasore, a dynamin inhibitor,^42,43^ significantly reduced dextran transport, consistent with previously reported indirect effects on membrane recycling and Rac1 activation (**Fig. 2D**).^44^ Imipramine, which impairs macropinocytosis by disrupting lipid raft domains and actin dynamics, significantly decreased permeability to FITC-dextran, a macromolecular probe known to enter via nonspecific fluid-phase pathways.^45^ Dynasore also reduced permeability of FITC-transferrin, which undergoes clathrin-mediated, receptor-driven uptake via TfR (**Fig. 2E**).^46^

Together, these data demonstrate that our mBBB model faithfully replicates key structural, molecular, and functional attributes of the human BBB and provides a tunable, imaging-compatible platform for studying nanoparticle interactions and transport under physiologically relevant conditions.

### 2.2 Diverse nanoparticle types exhibit distinct physical and biochemical properties

To investigate how physical and biochemical features influence nanoparticle transport across the BBB, three classes of nanoparticles were selected for comparative analysis: EVs derived from heterologous cell lines, synthetic liposomes, and polystyrene (PS) nanoplastics of two defined sizes (26 nm and 100 nm). Each nanoparticle type was characterized in detail for morphology, size distribution, labeling efficiency, and molecular composition (**Fig. S1**). These characterization steps ensured that the nanoparticle panel encompassed a broad range of sizes, structures, membrane compositions, and biological origins, thus providing the opportunity for an initial investigation of how specific features influence BBB transport outcomes in the mBBB system.

Synthetic liposomes were generated using microfluidic mixing. Formulations included PEGylated lipids and trace incorporation of a fluorescent lipid for membrane labeling. CryoEM imaging confirmed uniform vesicle morphology with a clear lipid bilayer and absence of internal structure (**Fig. S1A**). NTA measurements indicated a mode diameter of ∼105 nm (**Fig. S1E**). There is also presence of some smaller ∼15-20 nm vesicles observed in the EM images that are too small to detect by NTA. Heterologous EVs were isolated from two cell lines representing non-malignant (HEK 293T, or HEK)^47^ and brain-tropic malignant (MDA-MB-231-Br, or MDA-Br) origin.^48^ EVs were produced in CELLine bioreactors and purified by differential centrifugation followed by size exclusion chromatography (SEC). Cryogenic electron microscopy (cryoEM) confirmed the presence of intact vesicles with a lipid bilayer and internal granularity consistent with luminal cargo for both EV types (**Fig. S1B,C**). PS nanoplastics were selected to reflect environmentally relevant particle types found in human brain tissue.^6,49,50^ Two fluorescently labeled bead types, 26 nm (green) and 100 nm (red), were used. Scanning electron microscopy confirmed spherical morphology with minimal aggregation (**Fig. S1D**). Due to high index of refraction and poor scattering properties of the 26 nm beads, NTA was not used for concentration estimation of either NP bead populations; instead, nominal concentrations were calculated from stock solids content. Zeta potential measurements showed similar surface charge for both nanoplastics (**Table S1**).

Nanoparticle tracking analysis (NTA) revealed size distributions centered around ∼120 nm for HEK EVs and ∼140 nm for MDA-Br EVs (**Fig. S1F,G**), with overall particle concentrations in the 10⁹-10¹⁰ particles/mL range. Multiplexed immunocapture assays using tetraspanin markers CD9, CD63, and CD81 (**Fig. S1H**) confirmed EV identity and showed high marker co-localization for both HEK and MDA-Br EVs (**Fig. S1I,J**), consistent with established EV characterization guidelines.^51^ Dual-labeling with a lipophilic membrane dye (MemGlow 700) and cytosolic dye (CFSE) achieved >90% colocalization efficiency (**Fig. S1K**), allowing for discrimination between membrane-only and intact cargo-bearing vesicles in subsequent trafficking assays.

### 2.3 Nanoparticles exhibit source-dependent differences in BBB transport and accumulation

To evaluate how different nanoparticles interact with and traverse the BBB, the mBBB model was exposed to each of the five nanoparticle variants under standardized conditions. After 16 hours of incubation, confocal microscopy was used to assess nanoparticle accumulation, spatial distribution, and relative transport across the endothelial barrier into the ECM channels.

Striking differences in transport behavior were observed across nanoparticle types (**Fig. 3A-F**). HEK 293T-derived EVs showed the most prominent accumulation and penetration within the ECM. MDA-Br EVs also trafficked across the mBBB, though to a lesser extent. In contrast, bare liposomes and both 26 nm and 100 nm PS beads showed limited to negligible penetration beyond the endothelial monolayer, often appearing restricted to the luminal surface. Quantitative image analysis of positive fluorescence signal in the perivascular ECM region confirmed these observations (**Fig. 3G**). To probe the fate of EVs post-transit, dual-labeled EVs were used to assess co-localization of cytosolic and membrane signals. Both HEK and MDA-Br EVs demonstrated 5-10% co-localization in the ECM compartment, suggesting that a minority population of vesicles are intact following BBB transport (**Fig. 3H**).

**Figure 3.**
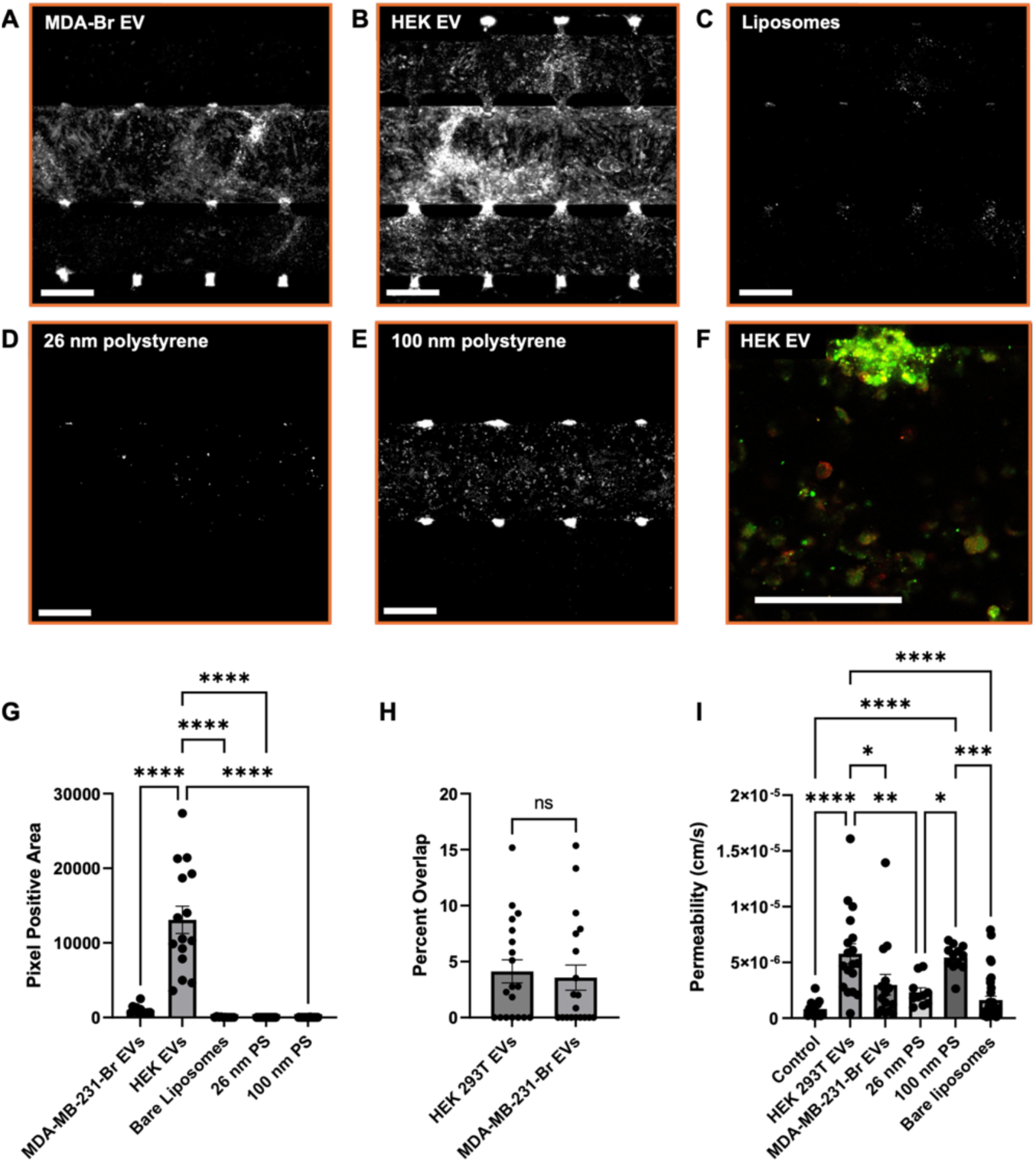
EVs exhibit enhanced transport across the mBBB compared to synthetic nanoparticles. (**A-E**) Representative confocal images showing nanoparticle accumulation across the mBBB. Fluorescent signal (grayscale) indicates EV or nanoparticle presence following 16-h incubation: **(A)** MDA-MB-231-Br EVs (MDA-Br EVs), **(B)** HEK 293T EVs (HEK EVs), **(C)** liposomes, **(D)** 26 nm PS beads, and **(E)** 100 nm PS beads. Magnification: 10x. Laser: 488 (EVs, 26 nm PS), 561 nm (100 nm PS), 640 nm (liposomes). Gain: 1.0. Exposure time: 2 µs/pixel. Scale bars: 100 µm. **(F)** High-magnification views of dual-labeled HEK EVs, showing colocalization of cytosolic (green) and membrane (red) dyes in the perivascular ECM. Magnification: 60x. Laser: 488, 640 nm. Gain: 1. Exposure time: 2 µs/pixel. Scale bar: 100 µm. **(G)** Total pixel-positive area across all ECM zones by particle type. HEK EVs (n=12) show the highest accumulation, significantly exceeding MDA EVs (n=15), bare liposomes (n=10), 26 nm PS (n=31), and 100 nm PS (n=30) by one-way ANOVA. **(H)** Percentage of dual-labeled EVs maintaining membrane/cargo co-localization after mBBB transit, indicating similar levels of intact transport for HEK (n=19) and MDA (n=19) EVs by unpaired t-test. **(I)** Post-treatment mBBB permeability to 70 kDa FITC-dextran reveals increased barrier disruption following exposure to HEK EVs (n=18) and 100 nm PS particles (n=12), but not to MDA EVs (n=15), liposomes (n=34) or 26 nm PS (n=10) when compared to control (n=17) by one-way ANOVA. (*) indicates p < 0.05, (**) indicates p < 0.01, (***) indicates p < 0.001, (****) indicates p < 0.0001, and (ns) indicates p > 0.05. Error bars indicated standard error in the mean (SEM). Replicate data are biological replicates. Sample size (n) indicates pores used in analyses.

These results demonstrate that biologically derived EVs exhibit superior BBB transport relative to synthetic particles and that protein expression profiles, particularly RMT ligands, may govern transcytosis efficiency. However, most EVs are retained near the endothelial interface, and only a fraction maintains structural integrity post-transport, posing challenges for therapeutic bioavailability.

### 2.4 BBB permeability is selectively disrupted by nanoparticle interaction at the endothelium

To assess whether nanoparticle exposure alters BBB integrity, we measured permeability to 70 kDa FITC-dextran following treatment with each nanoparticle type. This assay evaluates macromolecular passage across the endothelium and provides a functional readout of barrier disruption. Among the particles tested, HEK 293T EVs and 100 nm PS beads induced a 2– to 3-fold increase in dextran permeability relative to untreated controls (**Fig. 3I**). This suggests that both biologically active and synthetic nanoparticles can compromise BBB function. MDA-Br EVs, 26 nm nanoplastics, and liposomes did not impact dextran permeability.

Interestingly, barrier disruption did not correlate directly with transport efficiency. HEK EVs trafficked efficiently and increased permeability, while 100 nm PS beads, despite poor translocation, elicited a similar degree of disruption. This disparity suggests that BBB dysfunction may result from surface interactions or indirect signaling rather than bulk particle accumulation in the perivascular space.

Confocal imaging of the 100 nm PS condition revealed bright, punctate fluorescence localized near the endothelial interface, particularly at micropore regions where barrier cells contact one another (**Fig. 3E**). This accumulation likely reflects particle adhesion at the luminal surface rather than transcytosis. The observed increase in permeability in this context supports the hypothesis that surface-associated nanoplastics can disrupt endothelial function through membrane interaction, inflammatory signaling, or oxidative stress, even in the absence of deep penetration.

These results highlight the importance of evaluating both transport and toxicity when designing nanoparticles for brain delivery. While EVs may use endogenous trafficking machinery to enter the brain, synthetic particles can cause barrier dysfunction without entering the parenchyma, raising concerns for environmental and pharmaceutical nanoparticle safety.

### 2.5 Nanoparticle transport utilizes distinct cellular uptake mechanisms across the BBB

To elucidate the cellular pathways governing nanoparticle transcytosis across the BBB, we selectively inhibited two major endocytic mechanisms: dynamin-mediated endocytosis and macropinocytosis. Dynasore, a non-competitive inhibitor of dynamin GTPase activity, blocks clathrin– and caveolin-dependent vesicle scission.^42^ Imipramine disrupts lipid rafts and cytoskeletal remodeling, thereby inhibiting macropinosome formation.^43^ Each inhibitor was pre-incubated with the mBBB and maintained in solution during nanoparticle exposure. Dual-labeled EVs were used to separately quantify transport of membrane (lipophilic dye, MemGlow700) and luminal cargo (cytosolic dye, CFSE).

Dynasore significantly reduced both luminal cargo and membrane transport for HEK and MDA-Br EVs (**Fig 4A, B**), indicating broad reliance on dynamin-dependent pathways. Imipramine nearly abolished HEK EV transport (when tracking cytosolic dyes) but did not significantly reduce MDA-Br EV uptake (**Fig 4C, D**), suggesting that while macropinocytosis is essential for HEK EVs, MDA-Br EVs may engage additional, compensatory mechanisms. In contrast, synthetic particles behaved differently: Dynasore increased transport of liposomes, while Imipramine had little to no effect on any particles (**Fig 4E, F**). These results imply that synthetic nanoparticles bypass regulated uptake and may instead exploit nonspecific or destabilizing interactions with the endothelium.

**Figure 4.**
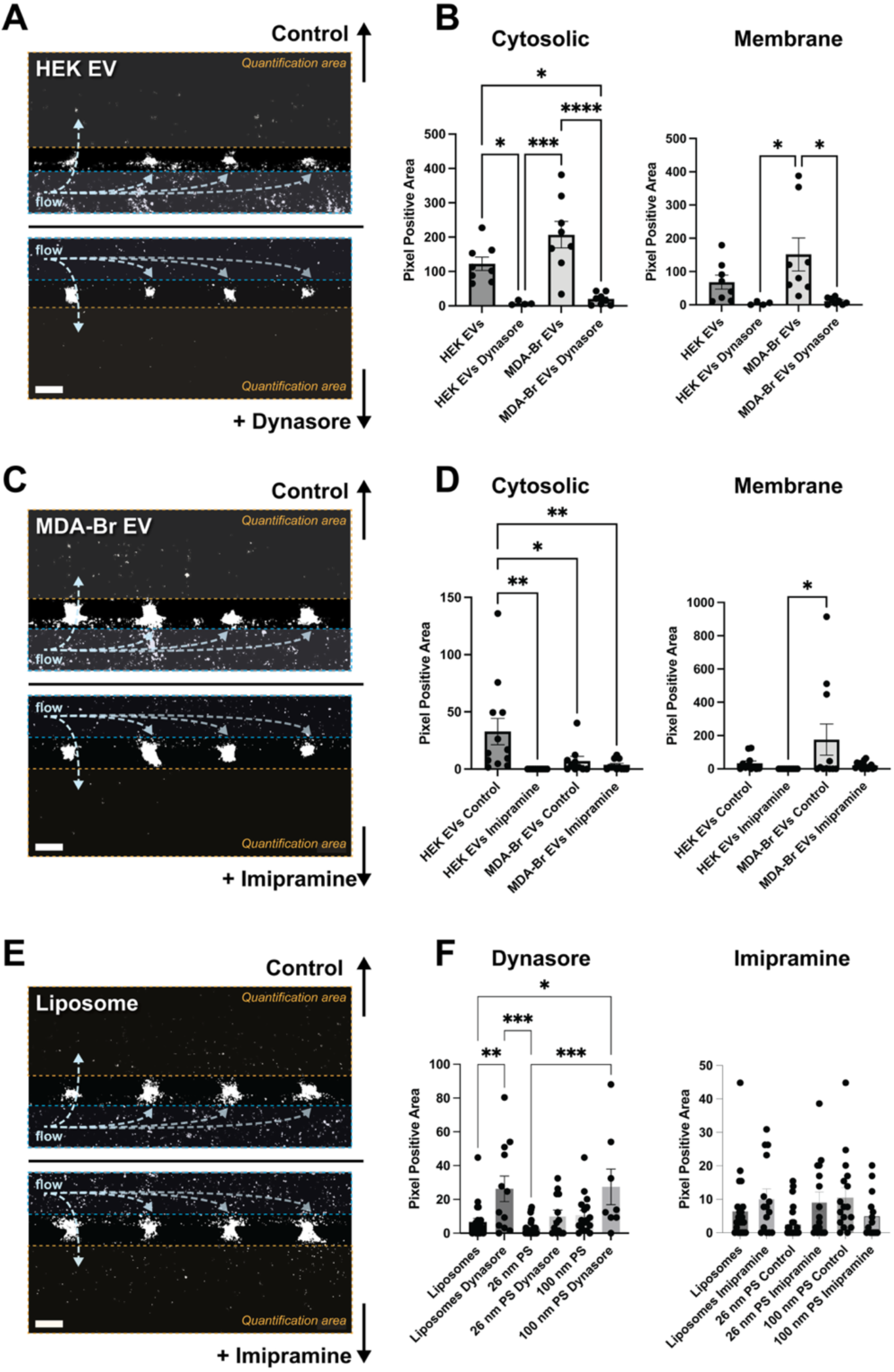
Pharmacological inhibition of endocytic pathways reveals distinct mechanisms of nanoparticle transport across the mBBB. (**A**) Representative confocal images of HEK EV accumulation in the ECM under control (top) and Dynasore-treated (bottom) conditions. **(B)** Quantification of cytosolic (CFSE, left) and membrane (MemGlow 700, right) signals shows that Dynasore significantly reduces cytosolic transport of both HEK and MDA-Br EVs. **(C)** Representative images of MDA-Br EVs under control (top) and Imipramine-treated (bottom) conditions. **(D)** Quantification indicates that Imipramine reduces HEK EV trafficking and partially reduces MDA-Br EV cytosolic and membrane transport. **(E)** Representative images of liposomes under control (top) and Imipramine-treated (bottom) conditions. **(F)** Quantification of synthetic nanoparticle trafficking (liposomes, 26 nm and 100 nm polystyrene beads) suggests that Dynasore significantly increases liposome transport, while Imipramine did not significantly decreases any particle transport. Nanoparticles were fluorescently labeled and imaged by confocal microscopy (10x objective; excitation 488 nm, 640 nm). Orange dashed boxes indicate quantification areas. Scale bars: 100 µm. All significance testing was performed using one-way ANOVA. (*) indicates p < 0.05, (**) indicates p < 0.01, (***) indicates p < 0.001, (****) indicates p < 0.0001, and (ns) indicates p > 0.05. Error bars indicated standard error in the mean (SEM). Replicate data are biological replicates. Sample size (n) indicates pores used in analyses.

Together, these findings demonstrate that nanoparticle transport across the BBB is not uniform, but instead governed by size, origin, and biochemical presentation. Biologically derived EVs utilize active, regulated mechanisms (including dynamin– and macropinocytosis-dependent pathways) that can be selectively modulated. In contrast, synthetic particles interact less specifically, and their trafficking may even increase upon inhibition of key cellular uptake systems, raising concerns about unintended permeability shifts induced by nanoparticle exposure or therapeutic co-administration.

## 3. Discussion

Nanoparticle trafficking across the BBB remains a central challenge in nanomedicine, neurobiology, and environmental toxicology. In this study, a horizontally oriented, spatially structured microfluidic BBB model was developed and validated to recapitulate key features of the human neurovascular unit. Composed of human endothelial cells, astrocytes, and pericytes embedded in a physiologically relevant ECM, this platform enables depth-resolved, high-content imaging and mechanistic interrogation of nanoparticle transport across the BBB. Using this model, we interrogated the transport and accumulation of three major classes of nanoparticles (EVs, liposomes, and nanoplastics) and were able to demonstrate that overall BBB transport is heterogeneous and depends on the source, size, and biochemical composition and presentation of the membrane. Furthermore, the model is simple enough in structure to be replicated by others and thus may be useful to explore a range of applications including mechanistic understanding of BBB transport as well as drug delivery design.

Studying BBB penetrance *in vivo* is only possible via expensive, low-throughput pre-clinical models that are confounded by interspecies differences, multi-organ clearance (via the reticuloendothelial system), or other complex interactions, such as with the immune system.^52^ While the latter are features of *in vivo* drug delivery in humans, it is difficult to tease apart these separate features in pre-clinical animal models. A variety of *in vitro* BBB models have been developed to evaluate BBB transport, including Transwell inserts, spheroid/organoid cultures, and microfluidic chips.^50,53–57^

While endothelial-only Transwell models are widely used due to their simplicity and compatibility with permeability assays, they often lack shear stress, 3D architecture, and neurovascular complexity, limiting their ability to fully recapitulate *in vivo* BBB behavior or resolve transcytosis mechanisms. For example, the role of astrocytes and pericytes in regulating receptor expression at the endothelial surface of the BBB is well established,^58^ and directly impacts nanoparticle transcytosis.^59–63^ Moreover, additional features of the human BBB are rarely considered, such as ECM stiffness, which impacts actin formation, tight junction localization, and phenotypic changes.^64–67^ Current mechanistic studies largely employ tissue culture plates or Transwell membranes with stiffnesses in the GPa range, orders of magnitude stiffer than brain ECM (0.2-1.2 kPa).^68,69^ Direct interactions between nanoparticles and the ECM can also dictate whether they survive transport across the BBB.^70,71^ Spheroid and organoid BBB models offer improved cytoarchitecture and tight junction expression,^72,73^ but lack perfusion and defined luminal compartments, complicating quantification of transport and accumulation of nanoparticles following BBB transport. More recently, microfluidic BBB chips, particularly vertical two-channel systems like the Emulate Brain Chip,^74^ have introduced flow and multi-lineage co-culture under physiologically relevant conditions. These platforms support more accurate measurement of drug or nanoparticle permeability but often suffer from limited imaging access due to the opaque membrane interface. Planar, horizontal microfluidic chips (Mimetas OrganoPlate^®^) have emerged to address these limitations by enabling real-time visualization of nanoparticle traversal and direct interaction with perivascular cells in 3D ECM environments and may prove useful in the development of *in vitro* 3D models of the BBB.^75^

The horizontal mBBB model described in this study builds on these advances, uniquely integrating perfusable endothelialized channels, embedded astrocytes and pericytes, and an optically accessible layout. Unlike vertical organ-on-chip designs, including conventional Transwell formats, this system permits simultaneous imaging of all barrier components, enabling spatially resolved analysis of nanoparticle localization, endocytic uptake, and transcytosis. The inclusion of flow and 3D ECM replicates the mechanical and biochemical cues of the *in vivo* BBB, supporting strong junctional expression and transporter functionality. ECM stiffness is known to affect tight junction formation, actin dynamics, and transcytotic activity, all of which may influence nanoparticle trafficking outcomes.

The mBBB’s open architecture also supports functional flexibility. Recovery of particles completing transcytosis allows for future proteomic or cargo-based analyses of BBB-penetrant subpopulations. The model’s compatibility with chemical inhibition, dual-color imaging, and fluorescence-based quantification enables mechanistic studies of entry pathways and downstream effects. To that end, the model was validated structurally by immunofluorescence for key junctional and transport proteins (ZO-1, Connexin-43, P-glycoprotein, transferrin receptor, LDLR, SR-BI) and functionally via low permeability to 70 kDa dextran and doxorubicin, consistent with *in vivo* BBB behavior.

Mechanistic perturbation using Dynasore and imipramine confirmed the engagement of dynamin-mediated and macropinocytic pathways, respectively, supporting the model’s capacity to emulate regulated endocytosis. Furthermore, additional cells present in the brain microenvironment including neurons and microglia can be easily incorporated in future iterations. Together, these features position the mBBB as a next-generation platform for mechanistic studies of nanoparticle trafficking and screening of brain-targeted nanotherapeutics.

We carried out a systematic comparison of five nanoparticles encompassing three major classes (two EV populations, synthetic liposomes, and two sizes of PS nanoplastics). Our results revealed clear distinctions in BBB transport. HEK 293T-derived EVs exhibited significantly greater trafficking across the mBBB than MDA-MB-231-Br EVs, liposomes, or PS beads. Moreover, mechanistic dissection using endocytosis inhibitors revealed that EVs rely on regulated uptake mechanisms: dynasore reduced cargo transport in both EV types, while imipramine abolished HEK EV trafficking entirely. Liposomes and PS beads, by contrast, had very low BBB penetrance under control conditions, and thus were unaffected or even increased in trafficking after inhibition, consistent with disruption of the BBB (**Fig. 3I**, 100 nm PS), and suggesting distinct, non-regulated entry routes or membrane destabilization effects. Although these data are correlative and obtained at the population level, they suggest that specific receptor-binding ligands contribute to enhanced transcytosis. Future studies may use receptor blockade or engineered EV surfaces to establish causality.

Dual-label tracking revealed that only 5-10% of detectable EVs survive intact passage across the mBBB. Fluorescent signal from both membrane and cytosolic dyes co-localized in only a minority of vesicles in the ECM, suggesting that most EVs are retained, degraded, or undergo partial fusion. Spatial analysis further showed that both EV populations tend to accumulate near the endothelial interface, with limited penetration into distal regions of the ECM. This observation may be an artifact of the device geometry, but also may reflect binding to ECM proteins. Regardless, it has potential implications for therapeutic EV use, as the effective dose reaching parenchymal targets may be lower than expected based on the total EV dose.

In contrast, liposomes and PS particles displayed poor BBB transport. Notably, 100 nm PS beads caused significant barrier disruption, as measured by increased dextran permeability, despite minimal transcytosis. Confocal imaging showed bead accumulation at the luminal endothelial surface, particularly near micropore regions, suggesting that physical or biochemical interaction with the cell membrane is sufficient to compromise barrier integrity. This is concerning for environmental nanoplastics, which may induce neurovascular dysfunction even without entering the brain parenchyma.

While we did not include a direct *in vivo* comparison in this study, the ranking of nanoparticle transport observed in our system aligns with prior reports of brain accumulation for EVs, liposomes, and nanoplastics.^50,55,70,76,77^ Recent studies using other BBB models have highlighted that nanoparticle and EV transport is governed by biochemical surface features, particularly receptor-targeting ligands. For example, nanoparticles functionalized with brain-targeting ligands, such as ApoE or transferrin, exhibit enhanced endothelial uptake and transcytosis compared to unmodified particles, despite comparable size and charge.^78^ Similarly, transferrin-conjugated antibodies demonstrated selective transcytosis in a perfused planar microfluidic BBB model.^79^ In therapeutic contexts, lipid nanoparticles (LNPs) carrying siRNA or mRNA payloads have shown poor BBB penetration unless modified with specific peptides or ligands.^80^

However, our study also challenges some assumptions in the field. While prior work often presumes that tumor-derived EVs exhibit enhanced BBB crossing due to aggressive tropism, we found that MDA-MB-231-Br EVs trafficked less efficiently than HEK EVs, suggesting that surface ligand composition (not cellular origin) may better predict transport behavior. Furthermore, we observed that 100 nm PS particles, which are often used as size-matched controls, disrupted the endothelial barrier despite minimal penetration. This finding aligns with recent work showing that aged plasma EVs increased BBB permeability and reduced tight junction expression in a dynamic microfluidic model, highlighting that toxicity and trafficking can be uncoupled.^81^ Together, these data reinforce that advanced BBB-on-chip systems are uniquely capable of dissecting nanoparticle and EV behavior across the barrier, and support the use of human-relevant, spatially resolved platforms like the mBBB to guide therapeutic nanocarrier design and EV mechanistic studies.

Several limitations of the current system merit discussion. First, hydrostatic pressure drives intermittent flow, which may not fully replicate the continuous shear stress of *in vivo* vasculature. Shear is known to influence tight junction stability and receptor expression, and future versions of the mBBB could incorporate perfusion systems to sustain physiologic flow. Second, the use of the HCMEC/D3 cell line, while convenient, exhibited limited claudin-5 expression. Incorporating primary or iPSC-derived endothelial cells could enhance model fidelity. Labeling strategies for EVs also present caveats. CFSE labeling relies on esterase activity and internal protein content, potentially biasing results against protein-poor or enzyme-deficient EVs. MemGlow 700 labels only the outer membrane, limiting the ability to track multivesicular or multilamellar EVs post-lysis. These biases may lead to underrepresentation of specific EV subpopulations. Orthogonal approaches, such as genetically encoded membrane tags or encapsulated reporters, could complement current methods. Functional validation of candidate ligands and analysis of post-transit subpopulations remain important next steps.

## 4. Conclusion

This study establishes a high-content, spatially structured mBBB model as a flexible platform for probing nanoparticle interactions with the human BBB. The system supports detailed analysis of transport mechanisms, barrier disruption, and structure-function relationships. By revealing that trafficking is governed primarily by membrane composition and surface ligands, not size or stiffness alone, these results provide a framework for rational nanoparticle design and screening. The mBBB thus addresses an unmet need in BBB research and offers a modular, human-relevant alternative for advancing both therapeutic delivery and neurotoxicity assessment.

## Methods

### Cell culture for mBBB

Human Cerebral Microvascular Endothelial Cells (HCMEC/D3) at passage 5 were cultured in endothelial cell media (ScienCell Research Laboratories, Cat. #1001) in flasks coated with 0.2% Gelatin. Prior to seeding, flasks were incubated with 0.2% Gelatin for >30 min at 37°C. Primary human brain vascular pericytes (HBVP; ScienCell Research Laboratories; Cat. #1200) at passage 4 were cultured on 20 μg mL^-1^ poly-L-lysine (PLL; ScienCell Research Laboratories; Cat. #0413) treated flasks. Primary human astrocytes (HA; ScienCell Research Laboratories; Cat. #1800) at passage 4 were cultured on 20 μg mL^-1^ poly-L-lysine (PLL; ScienCell Research Laboratories; Cat. #0413) treated flasks. Prior to seeding, flasks were incubated with 20 ug mL^-1^ PLL for >16 h at 37°C. PLL solutions were prepared by diluting with dPBS (−/−) (Thermo Fisher Scientific, Cat. #14190250). Cells were cultured for 4 days to reach 90% confluency prior to lifting for mBBB device seeding.

### Cell culture for EV production

The metastatic breast cancer cell line (MDA-MB-231-Br) was sub-cultured according to advised protocols using DMEM basal media (ATCC, Cat. #30-2002) supplemented with 10% fetal bovine serum (ATCC; Cat. #30-2020) and 1% penicillin/streptomycin (ATCC; Cat. #30-2300). The non-metastatic kidney line (HEK293T) was sub-cultured according to advised protocols using DMEM basal media (ATCC, Cat. #30-2002) supplemented with 10% fetal bovine serum (ATCC; Cat. #30-2020) and 1% penicillin/streptomycin (ATCC; Cat. #30-2300). Cells were cultured in T175 tissue culture flasks until 2.5×10^7^ cells could be collected and removed.

### EV production

CELLine AD 1000 bioreactors (EMD Millipore, Cat. #Z688045-3EA) were utilized to collect and concentrate EVs produced from each of the above outlined cell lines. Each bioreactor consists of a 15 mL cell compartment and 1 L media compartment separated by a 10 kDa cellulose semi-permeable membrane. 2.5×10^7^ of each cell type was diluted into 15 mL of supplemented media as mentioned previously and injected into the cell compartment of separate bioreactors (one cell type per bioreactor). 1 L of DMEM supplemented with 1% penicillin/streptomycin (Thermo Fisher Scientific; Cat. #15140163) was added to the media compartment. Twenty-four hours after cells were placed into the cell compartment, media in the cell compartment was replaced with serum EV depleted media. This media was prepared by obtaining media as mentioned above and removing serum EVs contributed by the fetal bovine serum. This was done by overnight centrifugation at 120,000 x g using an Optima LE-80K centrifuge. Prior to use, the EV depleted media was sterile filtered through a 0.22 μm pore size filter.

After 7 days, the media in the cell compartment was collected, the cell compartment was washed with 15 mL of dPBS −/− (Thermo Fisher Scientific; Cat. #14190250), and the media replaced with fresh EV depleted media. This protocol was repeated weekly for 12 weeks.

### EV isolation

EV containing media collected from the cell compartment was diluted 1:1 with 15 mL of dPBS −/− (Thermo Fisher Scientific; Cat. #14190250) to achieve a final volume of 30 mL. The EV solution was then sequentially centrifuged at 300 x g for 10 min, 2,000 x g for 15 min, and 10,000 x g for 30 min at 4°C. Between centrifuge cycles, the pellet was discarded. Following the 10,000 x g spin, the supernatant was placed into a fresh tube and stored at –80°C. Following 6 weeks of supernatant collection, all samples were further centrifuged at 120,000 x g for 70 min using an Optima LE-80K centrifuge. The supernatant was discarded, and the pellets were combined and suspended in 30 mL of 0.02 μm filtered dPBS −/− (Thermo Fisher Scientific; Cat. #14190250). The resuspended pellet was then further centrifuged for an additional 70 min at 120,000 x g. The pellet was then resuspended in 200 μL of 0.02 μm filtered dPBS −/− (Thermo Fisher Scientific; Cat. #14190250) and transferred to LoBind tubes (Fisher Scientific, Cat. #05-414-206). Purified EVs were then either used immediately or stored at –80°C until used.

### Single label EV preparation for trafficking and depth penetration

MDA-MB-231-Br and HEK 293T EVs were thawed from frozen and individually diluted to 75 μL using 0.02 μm filtered EBM. Diluted EVs were mixed at 1:1 with 75 μL of 8 μM CellTrace Far Red (Thermo Fisher; Cat. #C34572) diluted in EBM. EV-dye suspension was incubated at 37°C for 1 h. Free dye was removed using size exclusion chromatography (SEC). Concentrations were determined immediately prior to use in experimentation using nanoparticle tracking analysis.

### Dual label EV preparation for inhibition and survival studies

MDA-MB-231-Br and HEK 293T EVs were diluted to 5×10^12^ particles/mL respectively. EVs were then aliquoted to 5 μL and mixed with 70 μL of 40 μM Carboxyfluorescein Succinimidyl Ester or CFSE (ThermoFisher, Cat. #C34554) diluted with dPBS(−/−). EV-dye suspension was incubated at 37°C for 1 h. Then, 70 μL of 200 nM MemGlow 700 (Cytoskeleton; Cat. #MG05-02) was added and the resultant solution was incubated for another hour at room temperature. Free dye was removed using SEC. Concentrations were determined immediately prior to use in experimentation using nanoparticle tracking analysis.

### Generation of CF-NPs

CF-NPs were formulated using a 94:5:1 molar ratio of 18:1 (9-Cis) PC (DOPC) (Avanti Polar Lipids, Inc, Cat. # 850375C-200mg) to DSPE-PEG2000-MALEIMIDE (Avanti Polar Lipids, Inc, Cat. #880126C-25mg) to 18:1 PE CF (Avanti Polar Lipids, Inc, Cat. #810332C-1mg). Lipids were aliquoted to the specified molar ratio and the chloroform solvent was evaporated in a N_2_ atmosphere for 15 min. Lipid film was then reconstituted in 200 Proof ACS grade ethanol (Spectrum Chemicals & Laboratory Products, Cat. #E1028-1LTGL) to a final concentration of 50 mg/mL. Liposomes were then loaded into NanoAssemblr™ Spark™ cartridges (Cytiva Life Sciences, Cat. # NIS0013) requiring 48 μL of dPBS (−/−) in the collection well, 32 μL of dPBS in the aqueous well, and 16 μL of the lipid suspension in the organic well. Once all wells were filled, Spark™ cartridges were loaded onto the NanoAssmblr™ Spark™ instrument (Cytiva Life Sciences, Cat. # NIS0001) and formulation setting 3 was used. Liposomes were collected in a 0.5 μL LoBind centrifuge tube (Fisher Scientific, Cat. #05-414-206) and stored in a N_2_ or Ar rich environment overnight. Free floating lipid was removed using size exclusion chromatography. Concentrations were determined immediately prior to use in experimentation using nanoparticle tracking analysis.

### Size exclusion chromatography (SEC)

Following labeling of EVs or liposome synthesis, nanoparticles were isolated and purified from free dye or lipid. Nanoparticles were purified using Automatic Fraction Collectors (AFC) V2 with qEVsingle / 35 Gen2 (Izon, Cat. #ICS-35) on AFC V2 Firmware Version 1.2.2 to separate nanoparticles from free dye or lipid. Each isolation yielded nanoparticles in 0.8 mL of EBM using a default buffer volume of 0.87 mL. Following manufacturer guidelines, columns were flushed using 6 mL of 0.02 μm filtered EBM. Once 6 mL had been passed through the column, remaining media was removed and 150 µL of sample was added to the column. Once the sample had passed through the top frit into the column, 3 mL of fresh filtered EBM was added to the top to prevent column desiccation. EBM was filtered through a 20 nm Whatman® Anotop® 10 syringe filter (EMD Millipore Corporation, Cat. #WHA68091002) a 30 mL syringe (VWR International LLC, Cat. #302832). Isolated nanoparticles were then analyzed on NTA and used in experiments on the same day.

### Preparation of polystyrene beads

Fluoro-Max 26 nm green polystyrene beads (Thermo Scientific, Cat. #G25) and Fluoro-Max 100 nm red polystyrene beads (Thermo Scientific, Cat. #R100) were each vortexed individually to attain homogenous polystyrene suspensions. Particle count was calculated based on percent solids as reported by the manufacturer. When not in use, polystyrene beads are maintained at 4°C and covered in tin foil to avoid aggregation and photobleaching. When diluted for experimentation, 0.02 μm filtered EBM was used (Lonza, Cat. #CC-3121).

### Nanoparticle tracking analysis (NTA)

To obtain sizing and concentration data for EV samples, isolated EVs were diluted using 20 nm filtered dPBS (−/−) and analyzed using the NanoSight LM10 (Malvern Panalytical Ltd., UK). The LM10 has a 405 nm laser module, a sCMOS camera and an automated syringe pump (Harvard Bioscience, MA). Prior to injection of EVs, all components are pre-washed with 70% ethanol, MilliQ water, and dPBS (−/−). For accurate sizing and concentration measurements, EV samples were diluted to within 5×10^8^ and 1.5×10^9^ particles per mL. To acquire sizing and concentration information, three videos of EVs flowing through the imaging chamber of the LM10 were captured. Each of such videos were 30 s in duration and captured at a camera level of 11 or 12. Data analysis of these videos was performed using NanoSight NTA 3.4 with detection level set to 3. NTA figures shown are not dilution corrected.

### Cryo-Electron microscopy (CryoEM)

Copper lacey formvar grids (TedPella, Cat. #01883-F) were glow discharged for 1 min and 10 s at 30 mA negative polarity (Ted Pella, PELCO EasiGlow). Prior to glow discharge, grids were placed in a glass chamber with carbon side facing up and pressure was brought to 0.39 mBar for 10 s. Following glow discharge cleaning, grids were treated with 4 μL of poly-L-lysine (Sigma Aldrich, Cat. #P8920) to place a positive charge on the carbon side. Following a 2-min incubation at room temperature, the poly-L-lysine coating was wicked off and replaced with 4 μL of EV solution. Grids were immediately loaded into a Leica EM GP2 Plunge Freezer for blotting. After 2 min of EV incubation on the grids, blotting was performed for 6 s by pressing the carbon side of the grid to a wicking paper. Grids were subsequently flash frozen in liquid ethane and transferred to liquid nitrogen for storage until time of imaging. Imaging of the grids was performed on the ThermoFisher Glacios at 11,000 × and 45,000 × magnification. Post processing was performed using ImageJ in which a Gaussian Blur filter with a radius of 2 was applied to the images. Brightness and contrast of images was enhanced with sharpness increased by 36%, brightness decreased by 9%, and contrast increased by 55% for all images.

### Scanning Electron Microscopy (SEM)

Polystyrene beads were diluted in UltraPure water and 3 µL of each sample were dried on a silicon wafer. Samples were then sputtered with gold using a Denton Vacuum Desk II Cold Sputter/Etch Unit (Denton Vacuum Inc., Moorestown, NJ, USA) with a gold target at 200mTorr and 40 mA for 45-60 s dependent on PS bead size to mitigate charging effects. Sputter coatings were on the order of 10-30 angstroms. Sputter coating and scanning electron microscopy (SEM) were executed at the UC Davis Advanced Materials Characterization and Testing (AMCaT) laboratory. SEM was performed using a ThermoFisher Quattro S SEM (Thermo Fisher Scientific Inc., Waltham, MA, USA) equipped with a Schottky field electron gun. Secondary electrons were captured with an Everhart Thornley Detector (ETD). All images were captured with 15keV accelerating voltage and 0.22 nA current at a working distance of 5.0 mm (for 26 nm PS beads) and 10.0 mm (for 100 nm PS beads). All images were captured with a dwell time of 10.00 µs without drift correction.

### Tetraspanin immunocapture assay

Leprechaun “exosome human tetraspanin” kits (Unchained Labs) were stored in 4°C and chips were warmed to room temperature before use. Pre-warming chips help to prevent condensation which can affect chip binding. Chips were pre-scanned using provided protocols to obtain baseline measurements of chip backgrounds. Characterized EVs were diluted to 1×10^8^ particles/mL with 1× incubation solution (provided 10x incubation solution was diluted to 1x concentration with MilliQ water with 0.2% BSA).

This concentration was chosen to stay within the single particle regime (Mizenko et al., 2021). 50 µL of EV sample were prepared in the incubation solution to allow for excess when loading samples onto the chip. For sample incubation, chips were placed in the provided 24-well plate and 35 µL of diluted sample was pipetted onto the center of the chip. Care was made to ensure that sides of the chip were not contacting adjacent walls to prevent sample wicking. The 24-well plate was then sealed using manufacturer provided plastic adhesives, covered in aluminum foil and allowed to incubate overnight at RT. After incubation, 300 µL per chip of 1x blocking solution with 1 μg/mL of each antibody used. For manufacturer provided antibodies, 0.6 µL of antibody solution for each antibody was used. Chips were then washed with the ExoView CW1000 Plate Washer (Unchained Labs) on the CW-TETRA protocol. A total of 250 µL per chip of prepared blocking solution was added when prompted. Washed chips were then dried by wicking on a Kim Wipe. Once dried, chips were then transferred to the chuck and imaged using the ExoView R100 Automated Imager (Unchained Labs). Data was analyzed using ExoView Analysis Software (Unchained Labs). Thresholding for background was based on the mouse MIgG spot and set to limit particles detected to 100. Despite maintaining EVs in the single-particle regime, it is still possible that signal detected through the assay may not be single particle and positive pixel spots may represent groups of EVs rather than single EVs.

### mBBB microfabrication

mBBB devices were cast using the V031 device mold. V031 molds are placed into a petri dish and completely submerged in 100 grams of PDMS. PDMS was produced using a 10:1 ratio of Sylgard^TM^ 184 Silicone Elastomer Base to Sylgard^TM^ 184 Curing Agent (Ellsworth Adhesives; 184 SIL ELAST KIT 3.9KG). The Sylgard components are thoroughly mixed for >1 min prior to being placed into a desiccator to remove air bubbles introduced during mixing for 30 min. Following desiccation, the liquid PDMS precursor is transferred to the V031 molds and returned to the desiccator for an additional 30 min. Following the second desiccation, the V031 molds are cured in an oven at 65°C for >3 h. After curing over the molds, the PDMS blocks are cut and peeled off the molds. Inlet and outlet holes are punched into the ends of the created channels using a 13-gauge needle (Jensen Global, Cat. #JG13-0.5HPX) attached to a 5 mL BD Disposable Syringe with Luer-Lok™ Tip (Fisher Scientific, Cat. #1482945). PDMS blocks were flushed with N_2_ and vigorously washed under purified water. PDMS blocks were dried using N_2_ and baked for 5 min at 120°C. PDMS blocks and Epredia™ Richard-Allan Scientific™ Cover Glass (Fisher Scientific, Cat. #22-050-246) were plasma treated for 1 min before bonding. Devices were then baked for 15 min at 120°C prior to being stored until use. Tape was used to prevent dust or debris from entering channels prior to use. Before use, devices were sterilized via UV treatment for 30 min.

### Cell labeling for mBBB visualization

For visualization purposes, CellTracker dyes were used to label cells with different colors. Endothelial cells were labeled using CellTracker Green CMFDA, pericytes were labeled using CellTracker Orange CMRA, and astrocyte were labeled using CellTracker Deep Red (ThermoFisher, Cat. #C7025, #C34551, #C34585). Cells were labeled in tissue culture flasks prior to trypsinization by diluting 15 µL of stock solution into 10 mL of media. Cells were washed with dPBS (−/−) prior to addition of diluted dye. Cells were incubated for 30 min with the CellTracker dye at 37°C and 5% CO_2_. Cells were then washed with dPBS (−/−) and lifted using standard protocols.

### mBBB formation

Prior to adding cells into the devices, 200 μL tips were added to all outlet ports. Cells were then seeded into the microfluidic devices (**Figure S2**). Channels 2A and 2B were filled with HA and HVP suspended in a fibrin gel. Cells were concentrated to 2.5×10^7^ cells/mL at a 1:1 ratio and were resuspended in 10 mg/mL fibrinogen. The 10 mg/mL fibrinogen solution was formulated by mixing lyophilized fibrinogen (Sigma-Aldrich, Cat. #F8630-1G) in warmed dPBS (−/−) and incubating the solution in a 37°C water bath for >3 h. Once the HA and HVP cell suspension was made in fibrinogen, 15 μL was mixed with 0.9 μL of 50 U/mL thrombin. The thrombin solution was made by reconstituting lyophilized thrombin (Sigma-Aldrich, Cat. #T4648-10KU) in MilliQ water. The cell suspension was mixed with the thrombin solution 3 times and flowed through channels 2A and 2B from inlet to outlet until the levels of gel in the outlet and the inlet were equal. The pipette tip containing the gel was then ejected and left in the device. Devices were then placed into humidified 1000 μL pipette boxes and incubated at 37°C and 5% CO_2_ for 15 min. Devices were then removed from the incubator and channel 1 was filled with 10 μL of 50 μg/mL fibronectin (ScienCell Research Laboratories, Cat. #8248) diluted in dPBS (−/−). Devices were then incubated for an additional 15 min at 37°C and 5% CO_2_. During device incubation, the HCMEC cell suspension was prepared at a concentration of 25×10^6^ cells/mL in complete ECM media. Following incubation, channel 1 was filled with the HCMEC cell suspension. Channels 3A and 3B were then filled with complete ECM media. Devices were returned to the incubator for 2 h to facilitate cell adhesion.

Halfway through the 2-h incubation, devices were inverted to allow for effective coating of all surfaces. Following the 2-h incubation, all channels were flushed with EGM-2 (Lonza, Cat. #CC-4176). mBBBs were allowed to form for 3 days during which devices were fed by filling inlet tips of channel 1 with 100 μL of fresh EGM-2 and channels 3A+B with 20 μL of fresh EGM-2. EGM-2 media was mixed to be complete but without the VEGF supplement. Following three days of maturation, mBBB devices were analyzed for physiological relevance and downstream analysis.

### Immunostaining and image analysis

For the purposes of fixation, 200 μL of 10% formalin was added to the inlet side of channel 1 and allowed to flow through for 30 min. After 5 min of flow through, efflux from the outlet of channel 1 was aspirated to encourage flow of formalin. Following 30 min of fixation, formalin was removed from the inlet and outlet of channel 1 and 200 μL of 0.2% BSA in dPBS (−/−) was added. BSA solution was allowed to flow through the channel for 30 min with aspiration of efflux after the initial 5 min. After 30 min, the remaining BSA solution was removed and a fresh 200 μL of solution was added. Blocking with BSA was allowed to occur for an excess of 30 min up to 24 h. Staining of mBBB markers was performed for each of the following markers at the prescribed dilution factors as listed below in **Table 1**.

**Table 1:** Antibodies used to identify proteins expressed on endothelium including primaries, secondaries, and dilutions.

|  |  |  |
| --- | --- | --- |
| Antibody | Dilution factor/Concentration | Secondary antibody used |
| Recombinant Anti-Scavenging Receptor SR-BI antibody [EPR20190] (ab217318) | 1:100 | Goat-Anti Rabbit (Invitrogen A32733) |
| Recombinant Anti-LDL Receptor antibody [EP1553Y] (ab52818) | 1:100 | Goat-Anti Rabbit (Invitrogen A32733) |
| Recombinant Anti-Transferrin Receptor antibody [EPR20584] (ab214039) | 1:100 | Goat-Anti Rabbit (Invitrogen A32733) |
| Recombinant Anti-Connexin 43 / GJA1 antibody [EPR21153] (ab217676) | 1:100 | Goat-Anti Rabbit (Invitrogen A32733) |
| P-glycoprotein (human) monoclonal antibody [MRK16] (ALX-801-008) | 5 µg/mL | Donkey anti-Mouse (Invitrogen A31571) |
| ZO-1 antibody (ab2533977) (Cat# 71-1500) | 5 µg/mL | Goat-Anti Rabbit (Invitrogen A32733) |

Following a similar technique as described above, the primary antibody was incubated in the device for 2 h, followed by a 30-min washing step. The secondary antibody mixed with 5 μg/mL DAPI was incubated in the device for 1 h and again followed by a 30-min washing step. Devices were then imaged using confocal scanning laser microscopy. Post processing was performed in ImageJ to increase the brightness and contrast of images.

### Device allocation for experimental studies and blinding

Device allocation for experimental groups was performed blinded in which visual inspection of barrier formation was not used as a parameter for allocation between experimental and control groups. Further, similar strategies were employed for studies in which inhibitors were used for mechanistic manipulation. Investigator blinding was not performed for data collection but image acquisition was standardized to the 8 middle pores of the device in which all devices regardless of condition were imaged at the same frame. Analysis was blinded using an algorithm that treated each individual image in the same fashion.

### In vitro permeability assay against dextran, doxorubicin, and transferrin

Permeability analysis against dextran was performed following 3 days of mBBB formation, 4 days of mBBB formation, as well as following treatment with EVs on day 4. Permeability analysis against transferrin and doxorubicin was performed following 3 days of mBBB formation. 70 kDa FITC-dextran was reconstituted in MilliQ water to 10 mg/mL and further diluted with 0.02 μm filtered EBM to a final concentration of 100 μg/mL. Doxorubicin HCl (ThermoFisher; Cat. #J640000.MA) was diluted in dPBS (−/−) to a final concentration of 2 mM. FITC-Transferrin was reconstituted in MilliQ water and further diluted with 0.02 μm filtered EBM to a final concentration of 50 μg/mL. All precautions were taken to limit doxorubicin exposure as provided by the manufacturer and SDS. For permeability measurements, media in the mBBB was removed down to the PDMS block in the tips for channels 1, 3a, and 3b. 20 μL of either 100 μg/mL FITC-dextran, 50 μg/mL FITC-Transferrin, or 2 mM doxorubicin was then added to the inlet tip of channel 1 and imaged at 1.085 s per frame for 5 min. Videos were then imported to ImageJ. ROIs were drawn in channels 2a and 2b at the pore as well as one ROI in the middle of channel 1. Multi Measure was then used to measure the mean fluorescent intensity (I) value in the ROI across all 300 frames of the video. The data was then plotted against time. The slope (dI/dt) was determined in the linear range of the resultant figure and inserted into the following relationship^82^ to determine the permeability (cm/s) of the pore:

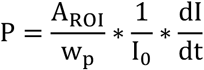

where I_0_ is the mean fluorescence intensity of the source (constant), A_ROI_ is the area (cm^2^) of the ROI, and w_p_ is the width (cm) of the pore. Note that mean fluorescence intensity is equivalent to the concentration of the fluorophore.

### Treatment of mBBB devices with EVs

EV treatment of devices was initiated on day 3 of mBBB formation. EVs were added to devices following pre-treatment of devices with either inhibitors, controls, or no pre-treatment. To dose mBBB devices with EVs, media was removed from the inlet and outlet tips of channels 1, 3a, and 3b. 100 μL of EVs diluted in 20 nm filtered EBM was added to the inlet tip of channel 1. The devices were imaged 16+ h following addition. Concentrations were dependent on yield following dye labeling and isolation steps. For initial trafficking studies, EVs were dosed at a concentration of 2×10^10^ particles/mL. For dynasore studies, EVs were dosed at a concentration of 4.3×10^9^ particles/mL. For imipramine studies, EVs were dosed at a concentration of 2.9×10^9^ particles/mL.

### Treatment of mBBB devices with bare liposomes

Liposome treatment of devices was initiated on day 3 of mBBB formation. Liposomes were added to devices following pre-treatment of devices with either inhibitors, controls, or no pre-treatment. Prior to treatment, liposomes were purified via SEC and counted using NTA. 100 μL of the liposome solution at 2×10^10^ particles/mL was added to mBBB devices and allowed to incubate for 16+ h in a humidified tip box in an incubator at 37°C and 5% CO_2_. Devices were imaged following incubation using confocal microscopy.

### Treatment of mBBB devices with polystyrene beads

Polystyrene bead treatment of devices was initiated on day 3 of mBBB formation. Beads were added to devices following pre-treatment of devices with either inhibitors, controls, or no pre-treatment. Fluoro-Max 26 nm green polystyrene beads (Thermo Scientific, Cat. #G25) and Fluoro-Max 100 nm red polystyrene beads (Thermo Scientific, Cat. #R100) were diluted to a theoretical concentration of 2×10^10^ particles/mL. 100 μL of each particle solution was added to independent mBBB devices and allowed to incubate for 16+ h in a humidified tip box in an incubator at 37°C and 5% CO_2_. Devices were imaged following incubation using confocal microscopy.

### Poisoning BBB uptake mechanisms

Dynasore (ABCAM Inc, Cat. # ab120192) and Imipramine (Sigma Aldrich, Cat. # I7379-5G) were used to inhibit dynamin associated transcytosis and micropinocytosis respectively. Both inhibitors were individually dissolved in DMSO at stock concentrations of 100 mM and 20 mM respectively. Working concentrations for each inhibitor were 50 µM and 10 µM, respectively. Inhibitors were diluted in the respective NP solutions ensuring that DMSO concentrations remained below 0.05%. Inhibition with Dynasore required a 30-min pre-incubation at 37°C and 5% CO_2_ prior to NP addition. Inhibition with Imipramine required a 60-min pre-incubation at 37°C and 5% CO_2_ prior to NP addition. Inhibitors were kept in solution with NPs to continue effective inhibition of each respective uptake pathway. Prior to addition of inhibitors, mBBB devices were washed with 200 μL of EBM. Pre-incubation was performed with 100 μL of inhibitor solution. NP treatments were performed using 100 µL of NP-inhibitor solution.

### Proteomics

Bulk EV protein proteomics was performed by the UC Davis Proteomics Core, Protein concentrations of EVs were analyzed using a Pierce micro-BCA assay kit (ThermoFisher, Cat. #23235). Prior to BCA analysis, EVs were lysed using RIPA lysis and extraction buffer (ThermoFisher; Cat. #89900). For each 15 μL of EV sample used, 8 μL of RIPA buffer was added and the solution was vortexed. EV-lysis solution was incubated on ice for 30 min prior to BCA assay. EV samples were provided undigested to the core for digestion and analysis. EVs were digested on S-Trap.

### Inclusion and exclusion criteria

Due to noticeable differences between the top and bottom pores of devices (in which top pores were more physiologically relevant in terms of permeability to small molecules), device bottoms were excluded from all studies with only device tops used for analytical purposes. Further, statistical outlier exclusions were employed as described below.

### Statistical Methods

All experimental values are expressed using mean and spread using standard error of the mean (SEM). Statistical significance between groups was evaluated using Student’s t-test and one-way analysis of variance (ANOVA) followed by Tukey’s post hoc test. Outliers were identified using ROUT with a threshold of 1%. GraphPad Prism version 10.4.2 was used for all data curation and statistical analysis. Statistical significance was evaluated at p values < 0.05. Error bars indicate significance differences, the absence of error bars between groups indicates a lack of significance between those groups. (*) indicates p < 0.05, (**) indicates p < 0.01, (***) indicates p < 0.001, (****) indicates p < 0.0001, and (ns) indicates p > 0.05.

## Author Contributions

**Bryan B. Nguyen**: Conceptualization (equal); data curation (equal); formal analysis (equal); investigation (equal); methodology (equal); writing – original draft preparation (equal); writing – reviewing and editing (equal). **Neona M. Lowe**: Data curation (supporting); methodology (supporting). **Sophia Kellogg**: Investigation (supporting). **Kuan-Wei Huang**: Investigation (supporting); Formal analysis (equal); methodology. **Hannah O’Toole**: Investigation (supporting); Formal analysis (supporting); methodology. **Elizabeth Hale**: Investigation (supporting); Formal analysis (supporting); methodology. **Venktesh S. Shirure:** Formal analysis (supporting); methodology. **Bhupinder S. Shergill:** Formal analysis (equal); methodology. **Steven C. George**: Conceptualization (equal); data curation (equal); formal analysis (equal); funding acquisition (equal); investigation (equal); methodology (equal); project administration (equal); writing – reviewing and editing (equal). **Randy P. Carney**: Conceptualization (equal); data curation (equal); formal analysis (equal); funding acquisition (equal); investigation (equal); methodology (equal); project administration (equal); writing – original draft preparation (equal); writing – reviewing and editing (equal).

## Acknowledgements

The authors acknowledge support from the National Institutes of Health (NIH), including R01CA241666, R01EB033389 and R01EB034279. NL was supported by a National Institute of Heart, Lung, and Blood Institution (NHLBI) funded training grant program (T32HL007013). BN was supported by T32GM144303. H.J.O. acknowledges T32GM136597. EJH was supported by a National Institute of Environmental Health Sciences funded training program in Environmental Health Sciences (T32 ES007059). This project was also supported by University of California Davis Biological Electron Microscopy (BioEM) Facility, which is supported by user fees, the Department of Molecular and Cellular Biology, the College of Biosciences, the Office of Research and the Provost’s Office. The Technical Director, Dr. Fei Guo, is supported by discretionary funds provided by MCB. The K3 and DED detectors were purchased from funding support provided by the Department of Molecular and Cellular Biology, College of Biological Sciences and grant support provided by R00-GM080249 (J. Al-Bassam). This work was also supported by the College of Engineering Next Level Research Seed Funding (UC Davis) and the School of Medicine Cultivating Team Science Program (UC Davis). The funders had no roles in the study design, data collection, data analysis, decision to publish, or preparation of the manuscript. The content of this work is solely that of the authors and does not necessarily represent the views of the funding parties. Components of figures in this work were created using or inspired by BioRender.com.

## Supporting Information

**Figure S1.**
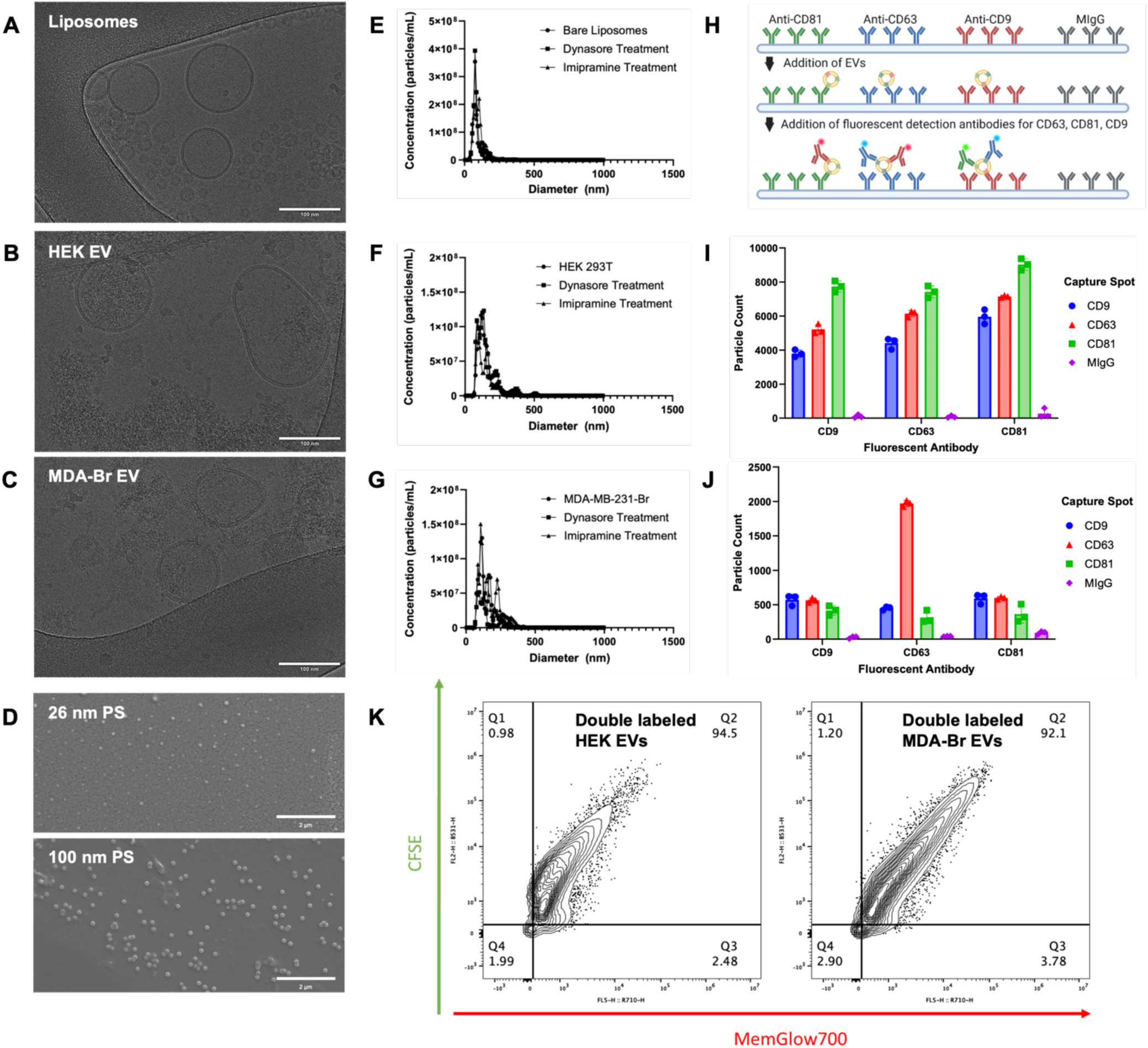
Characterization of diverse nanoparticle types used for BBB transport studies. (**A-C**) CryoEM images of representative nanoparticle types: **(A)** synthetic bare liposomes, **(B)** HEK 293T-derived EVs (or HEK EVs), and **(C)** MDA-MB-231-Br EVs (or MDA-Br EVs). All particles display characteristic lipid bilayers; EVs show internal granularity, while liposomes lack luminal structure. Magnification: 42,000x. Exposure time: 2 s. Gain: 40 electrons/angrstrom^2^. Scale bars: 100 nm. **(D)** Scanning electron microscopy (SEM) images of PS nanoplastics: *(top)* 26 nm beads and *(bottom)* 100 nm beads. Both exhibit spherical morphology and minimal aggregation. Magnification: 20,000x. Scale bars: 2 μm. **(E-G)** Nanoparticle tracking analysis (NTA) data for **(E)** synthetic bare liposomes, **(F)** HEK EVs, and **(G)** MDA-Br EVs. These data show particle size distributions and concentrations for each nanoparticle type, confirming expected dispersity and consistent batch characteristics. Control Dynasore and imipramine treatments do not significantly alter particle sizing or concentration across any particle type. **(H)** Schematic of ExoView-based tetraspanin immunocapture assay used to confirm EV identity via multiplexed labeling of CD9, CD63, and CD81. **(I-J)** Bar graphs show EV counts per tetraspanin and control (MIgG) capture spots for each fluorescent antibody channel for **(I)** HEK EVs and **(J)** MDA-Br EVs. **(K)** Flow cytometric analysis of EV double-labeling efficiency using a lipophilic membrane dye (MemGlow700) and cytosolic dye (CFSE), showing >90% colocalization for both HEK 293T (left) and MDA-Br (right) EVs. (*) indicates p < 0.05, (**) indicates p < 0.01, (***) indicates p < 0.001, (****) indicates p < 0.0001, and (ns) indicates p > 0.05. Error bars indicated standard error in the mean (SEM).

**Figure S2.**
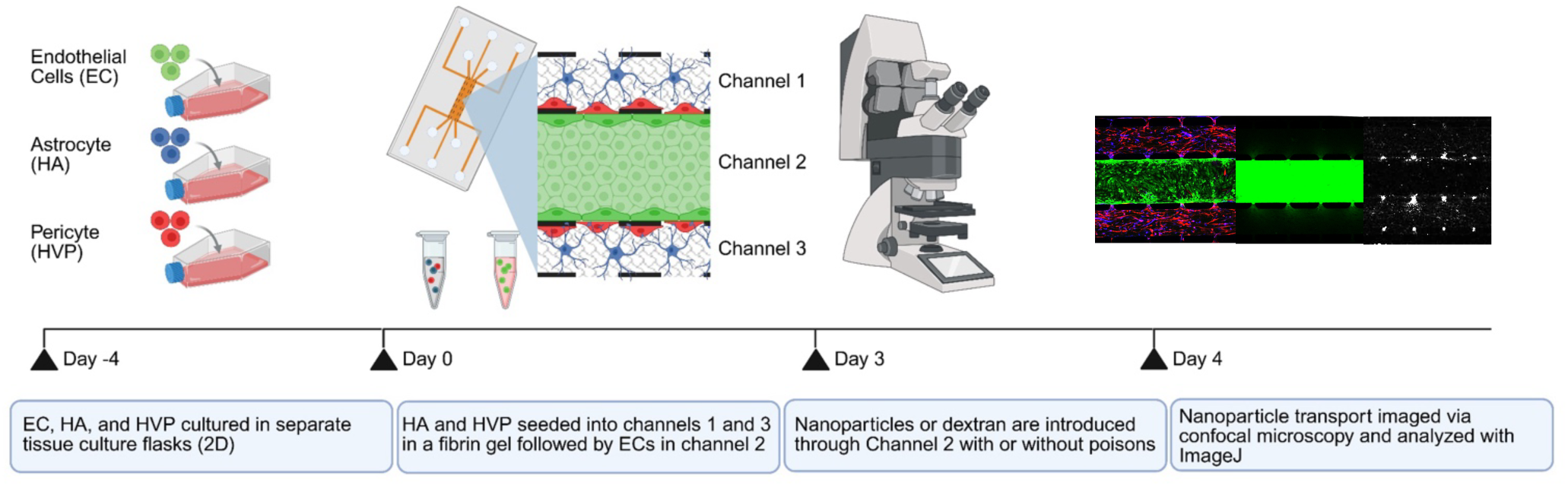
Experimental workflow for establishing and interrogating the microfluidic BBB model (mBBB). Schematic overview of the four-day experimental timeline used to establish and assay the mBBB. Endothelial cells (ECs), astrocytes (HAs), and pericytes (HVPs) are cultured separately in 2D conditions prior to introduction into the device. On Day 0, HAs and HVPs are suspended in fibrin gel and seeded into side channels (channels 1 and 3), followed by EC seeding into the central channel (channel 2). By Day 3, a physiologically structured BBB is formed, at which point fluorescent tracers or nanoparticles (with or without pathway-specific inhibitors) are introduced into the vascular channel. On Day 4, confocal microscopy is used to visualize and quantify nanoparticle trafficking across the mBBB. Imaging data are analyzed using ImageJ to assess transport efficiency, barrier integrity, and spatial distribution.

**Table S1.** Zeta potential measurements of polystyrene beads show similar surface charge for both nanoplastics.

|  |  |  |
| --- | --- | --- |
| Sample | Buffer | Zeta Potential (mV) |
| Fluoro-Max 26 nm green polystyrene beads | HEPES + 1% Tween 20 | -15.3 |
|  |  | -15.7 |
|  |  | -16.3 |
| Fluoro-Max 100 nm red polystyrene beads | HEPES + 1% Tween 20 | -17.9 |
|  |  | -18.4 |
|  |  | -16.4 |

